# Gender and speech material effects on the long-term average speech spectrum, including at extended high frequencies

**DOI:** 10.1101/2024.08.21.607381

**Authors:** Vahid Delaram, Margaret K. Miller, Rohit M. Ananthanarayana, Allison Trine, Emily Buss, G. Christopher Stecker, Brian B. Monson

**Affiliations:** Department of Speech and Hearing Science, University of Illinois Urbana-Champaign, Champaign, Illinois, 61820, USA; Boys Town National Research Hospital, Center for Hearing Research, Omaha, Nebraska, 68131, USA; Department of Otolaryngology/HNS, University of North Carolina at Chapel Hill, Chapel Hill, North Carolina, 27599, USA; Neuroscience Program, University of Illinois Urbana-Champaign, Champaign, Illinois, 61820, USA; Department of Biomedical and Translational Sciences, Carle Illinois College of Medicine, University of Illinois Urbana-Champaign, Champaign, Illinois, 61820, USA

## Abstract

Gender and language effects on the long-term average speech spectrum (LTASS) have been reported, but typically using recordings that were bandlimited and/or failed to accurately capture extended high frequencies (EHFs). Accurate characterization of the full-band LTASS is warranted given recent data on the contribution of EHFs to speech perception. The present study characterized the LTASS for high-fidelity, anechoic recordings of males and females producing Bamford-Kowal-Bench (BKB) sentences, digits, and unscripted narratives. Gender had an effect on spectral levels at both ends of the spectrum: males had higher levels than females below approximately 160 Hz, owing to lower fundamental frequencies; females had ∼4 dB higher levels at EHFs, but this effect was dependent on speech material. Gender differences were also observed at ∼300 Hz, and between 800-1000 Hz, as previously reported. Despite differences in phonetic content, there were only small, gender-dependent differences in EHF levels across speech materials. EHF levels were highly correlated across materials, indicating relative consistency within talkers. Our findings suggest that LTASS levels at EHFs are influenced primarily by talker and gender, highlighting the need for future research to assess whether EHF cues are more audible for female speech than for male speech.

## I. INTRODUCTION

Speech has audible acoustic energy at frequencies that span the frequency range of human hearing, including frequencies above 8 kHz and even above 13 kHz (Monson et al., 2014b; Monson and Caravello, 2019). The use of high-fidelity speech recordings that include energy at frequencies above 8 kHz, defined as extended high frequencies (EHFs), has not been standard practice for testing speech perception, particularly for speech materials that are widely used for clinical assessments (Monson and Buss, 2022). This may be due to technological constraints and/or a longstanding belief that EHFs have a negligible contribution to speech perception (Monson et al., 2014a). Recent studies have documented benefits of EHF cues for speech perception. For example, by low-pass filtering target speech and/or maskers at 8 kHz and comparing to full-band conditions, it has been demonstrated that EHF cues in target speech improve speech-in-noise recognition (Flaherty et al., 2021; Hunter et al., 2020; Monson et al., 2023; Motlagh Zadeh et al., 2019; Polspoel et al., 2022; Trine and Monson, 2020). Low-pass filtering speech at 8 kHz degraded talker head-orientation discrimination (Monson et al., 2019) and speech localization in elevation (Best et al., 2005). Removal of bands that included at least some energy above 8 kHz also led to degraded speech clarity (Füllgrabe et al., 2010) and word learning and recall (McCreery and Stelmachowicz, 2013; Pittman, 2008).

The utility of speech EHF cues for normal-hearing listeners depends on the audibility of EHFs in speech (Monson et al., 2023), which in turn depends on speech spectral levels at EHFs. Here we were motivated, in part, by a desire to assess factors that might influence EHF spectral levels in speech, including gender and speech material, to guide stimulus selection for speech recognition tasks. One challenge in attempting to study EHF cues is the lack of high-fidelity speech materials that faithfully represent realistic EHF spectral levels. For example, Monson and Buss (2022) analyzed the spectra of several popular speech corpora that are widely used both clinically and for speech perception research. Although some of these speech corpora include EHF content, many others were recorded decades ago with low sampling rates and/or microphones with limited frequency responses, leading to reduction or omission of EHFs. Thus, many of these materials fail to provide the full spectral range of acoustic cues available in real-world speech signals. Here we used a new corpus of high-fidelity, anechoic speech materials to investigate the effect of gender and speech material on the long-term average speech spectrum (LTASS), including at EHFs.

The LTASS is a well-established technique for evaluating the spectral content of speech signals. It is calculated by computing the frame-by-frame spectra (i.e., fast Fourier transforms) for contiguous, brief time frames across the length of a speech signal, and then computing the average of these spectra. An alternative LTASS method is to filter the time-domain speech signal into contiguous bands (e.g., third-octave bands) and calculate the root-mean-squared amplitude of each of the filtered signals, providing a time-averaged amplitude value for each frequency band. The LTASS has the advantage of capturing the time-averaged acoustic energy across frequency, which can be a useful way to characterize long speech signals. However, it also has disadvantages of tending to be dominated by phonemes that are long in duration (e.g., vowels) and failing to represent dynamic changes in the distribution of spectral energy across time.

There are several factors that can influence the LTASS of a speech recording, such as recording quality (Monson and Buss, 2022), emotion and prosody (Williams and Stevens, 2005), and vocal effort (Monson et al., 2012b; Sundberg and Nordenberg, 2006). Past studies have investigated the effects of gender and language on speech spectral levels using the LTASS. Although some of these studies focused on the frequency range below 8 kHz (Byrne, 1977; Olsen et al., 1987), others included EHFs in their analyses. These studies are reviewed here. The recording setup, including microphone distance and location, can also affect the LTASS, especially at EHFs. For example, the practice of using an off-axis microphone placement reduces EHF band levels because of increased directionality of speech radiation at EHFs (Kocon and Monson, 2018; Monson et al., 2012a). For this reason, these relevant recording details are reported for each study when they were available.

Cox and Moore (1988) calculated spectra up to 12.5 kHz for American English with 30 male and 30 female subjects. They recorded two minutes of continuous speech with a microphone (Bruel & Kjaer, type 4145) located at 0° azimuth and a 30-cm distance in a double-walled sound treated room. The male-female comparison revealed significant differences in third-octave band levels at both ends of the spectrum: below 200 Hz (higher levels for males) and at 8, 10, and 12.5 kHz (higher levels for females). They also reported gender differences in the 315-Hz band (higher for males), and the 800-Hz and 1000-Hz bands (higher for females).

Byrne et al. (1994) analyzed the LTASS for a wider frequency range up to 16 kHz. Cassette recordings of subjects with different native languages including American English were made while subjects read a passage from a storybook in their own language. Recordings were made in several countries in anechoic rooms whenever possible. In each location, a custom-made microphone unit was positioned 20 cm away from the talker at 45° azimuth. At least 10 male and 10 female subjects were recorded for each language, although the gender ratio for each language was not specified. Mean spectral levels showed females had about 0.3 to 5.9 dB higher levels than males (depending on the language) for the frequency range 6.3-12 kHz, and males had higher levels for frequencies <200 Hz. No statistical analysis was conducted to test these gender differences. That study found a significant effect of language on the LTASS, including differences observed at frequencies >6.3 kHz, with some evidence for an effect of dialect within language. Notably, the microphones used in that study had a frequency response that was flat only up to 10 kHz. Reported levels for frequencies above this limit were extrapolated and are therefore not entirely reliable. Additionally, the 45° microphone placement likely affected EHF levels as described earlier. The mean attenuation in EHF level at 45° (relative to 0°) is ∼3 dB (Monson et al., 2012a). However, there are large individual differences across talkers in EHF directionality that result in attenuations that can vary by as much as 7 dB at 45° (Monson and Ananthanarayana, 2023). Thus, the use of a 45° microphone placement conflated this additional source of individual variability with the individual variability in EHF level that would be observed with recordings made at 0°. This is also true for lower frequencies in the LTASS, although individual differences in directionality at lower frequencies are not as dramatic as those seen at EHFs (Monson and Ananthanarayana, 2023).

Moore et al. (2008) examined the LTASS at high frequencies for British English. That study included 8 female and 9 male subjects who read a prose passage at normal conversational level inside a double-walled sound treated room. Recordings were made with the microphone (Sennheiser MKH 40 P 48 U3) positioned 30 cm away and 15 cm below the talker’s mouth.

Microphone recording angle was not reported in this study. No statistical analysis was conducted to compare male and female levels, but the mean spectral levels suggested higher levels for females at frequencies >6300 Hz and higher levels for males at frequencies <200 Hz, as others have reported. Results of Moore et al. (2008) indicate 1-6 dB lower EHF levels compared to Byrne et al. (1994) when spectra were equated in overall level. The authors suggested that this EHF difference could be due to poor microphone frequency response and higher equivalent input noise in the Byrne et al. (1994) study.

Monson et al. (2012b) investigated the characteristics of high- and extended high-frequency energy in long-term average spectra of singing and speech for three production levels (soft, normal, and loud). That study included 8 female and 7 male subjects, and speech material consisted of six-syllable, low-predictability American English phrases. A Class I microphone was positioned at a 60-cm distance from the talker’s mouth at 0° azimuth and at the level of the mouth in an anechoic chamber. Monson et al. (2012b) reported an increase in high-frequency energy levels of speech with an increase in production level, and overall higher levels in singing voice compared to speech in soft and normal production levels. EHF third-octave band levels for females were significantly higher than for males, with mean differences between groups ranging from 4 dB (at 20 kHz) to 6.6 dB (at 10 kHz). This trend of higher female levels at EHFs was consistent when production level varied between speaking soft, normal, and loud. Third-octave band levels at EHFs calculated for normal speech were >5 dB higher than those reported by Byrne et al. (1994). The reason for this difference is not clear, although possibilities include the different phonetic content of the speech material, the quality and location of the microphones, room acoustic effects, and/or the inclusion of only vocally trained subjects. Furthermore, it is not clear whether the gender difference reported by Monson et al. (2012b) is generalizable because that study had a small sample size and included only young (all but one under the age of 30), vocally trained singers from the western United States.

The goal of the present study was to investigate the effect of talker gender and speech material on speech spectral levels, particularly at EHFs, using an anechoic multi-directional corpus of American English speech materials captured with high fidelity in the EHF range (Monson et al., 2022). One question we sought to address was to what extent materials with differing phonetic content might explain spectral differences seen across past LTASS studies. To assess this, we used different speech materials uttered by the same set of talkers. The speech materials consisted of Bamford-Kowal-Bench lists (BKBs; Bench et al., 1979), digits 0-10, and unscripted narrative speech. Based on the previous studies, we hypothesized that EHF spectral levels in the corpus would be higher for female speech than for male speech. As voiceless fricatives are known to have substantial EHF energy (Monson et al., 2012b), and five out of the eleven digits contain voiceless fricatives, we also expected higher EHF levels in digits relative to BKBs and unscripted narratives.

## II. METHODS

Acoustical analyses made use of the corpus described in detail by Monson et al. (2022). Methods for elicitation and capture of the speech materials are briefly summarized here.

### A. Speech material

The corpus consists of speech samples collected from 30 native speakers of American English (15 female, 15 male) with an age range of 21.3 to 60.5 years (mean 33.6) recruited from the community surrounding Omaha, NE. Gender was self-reported and corresponded with the participants’ biological sex. Based on their geographical location of residence before age 18, subjects represented various dialectal groups, including West, North, Midland, South, Mid-Atlantic, and New-England (Labov et al., 2008, pp. 119–151). Six subjects had a history of voice training. Participants’ self-identified racial backgrounds were: Asian (*n*=1), Black (*n*=1), Native American/Indian (*n*=1), White (*n*=24), more than one race (*n*=1), and undisclosed *(n*=2).

The speech samples were captured under anechoic conditions using a free-field Class 1 condenser microphone with a flat frequency response up to 20 kHz (B&K type 4189). The microphone was located 1 m directly in front of the seated talker (0° azimuth and equal height). The acoustic noise floor was < 15 dB SPL form 160-20,000 Hz. Recordings were stored as 24-bit uncompressed audio at a sampling rate of 48 kHz. The corpus consists of multi-directional recordings spanning 0° to 180° azimuth with 11.25° angular resolution, but only the 0° recordings were analyzed here.

Three sets of speech materials were extracted from the corpus for acoustical analysis. The first was unscripted conversational speech, in the form of a narrative recorded prior to the other materials. The second set consisted of spoken digits “zero” through “ten” recorded after the narratives. Digit productions were elicited following auditory presentation of a male template recording via an overhead loudspeaker. The third set consisted of 64 BKB sentences (lists 1-4; Bench et al., 1979) recorded after the digits and elicited in the same manner, using a female template recording. On average, the duration of the different materials was 2.5 min for narratives, 9 s for concatenated digits, and 1.8 min for concatenated BKB sentences. Due to equipment malfunction during recording, narratives were not captured for one male and one female talker. Additional details regarding the generation of the corpus, the recordings themselves, along with ancillary data, can be obtained from an open repository (Monson et al., 2022).

### B. Acoustical analysis

All audio was high-pass filtered using a fourth-order Butterworth filter at 70 Hz to reduce low-frequency noise. To calculate the LTASS for each subject and each speech material, a single audio file was created for each subject and speech material by concatenating individual sentence (BKBs) or digit recordings, with 50-ms cosine ramps to avoid discontinuities at sentence boundaries. For narratives, silent intervals over 200 ms were manually trimmed to be ∼200 ms. This analysis used recordings from the microphone at 0°, directly in front of the seated talker.

The root-mean-squared sound pressure level (SPL) was calculated for each speech material for each talker. The mean values at 1 m distance were 62.1 dB SPL for BKBs (range: 55.6-66.2 dB SPL), 64.6 dB SPL for digits (range: 60.8-67.7 dB SPL), and 63.1 dB SPL for narrative recordings (range: 54.2-69.1 dB SPL). For acoustical analysis, all recordings were scaled to 65 dB SPL, as done by Moore et al. (2008), which was in the range of sound levels measured here. The long-term speech spectra for each speech material and talker were calculated using a 2048-point fast-Fourier transform (FFT), resulting in a frame length of 42.7 ms and spectral resolution of 23 Hz, using a Hanning window and 50% overlap. An equivalent rectangular bandwidth (ERB; Glasberg and Moore, 1990) LTASS was calculated from the FFT LTASS by filtering and summing energy in the frequency domain. The ERB LTASS was calculated using 1-ERB-wide analysis bands (at 3-dB down points) with cosine-shaped filters centered at ERB numbers 3-41 with 50% overlap, similar to McDermott and Simoncelli (2011). To compare with previous studies, a third-octave band LTASS was also calculated from the FFT LTASS, using bandpass rectangular filters centered at third-octave frequencies from 79-20159 Hz with no overlap, similar to Monson et al. (2012b).

To analyze the phonetic content of the different speech materials, scripts (for BKBs and digits) and transcripts (for narratives) were converted to phonetic representations using the online tool toPhonetics.com. Narrative recordings were transcribed using the closed caption feature of Zoom software (Version 6.0.11) and then were manually corrected for transcription errors. We also analyzed the phonetic content of the script used in Monson et al. (2012) and the script of the Rainbow passage (Fairbanks, 1960, pp. 124–129) for comparison. The number of times each phoneme occurred in each recording was calculated using MATLAB (The MathWorks Inc., Natick, MA). The phoneme density was calculated by dividing the number of repetitions by the total number of phonemes.

### C. Statistical analysis

Data were analyzed using R (R Core Team, 2023). Linear mixed-effects models, with a random intercept for subject, were computed using the lme4 library (Bates et al., 2015). Models tested the influence of gender and speech material on EHF levels using ERB bands 34-41 (center frequencies [CFs] 8706 through 18769 Hz). EHF band number (i.e., frequency) was treated as a continuous variable because of an observed linear trend between frequency and band level in the EHF range. Band 34 was treated as the intercept point in the model. Interaction terms were initially included in the model but higher-level interactions that did not approach significance (*p*>0.15) were removed from the model. This criterion (*p*>0.15) reflects a conservative approach to interaction term removal, preventing removal of interaction terms that may be appreciably influencing model estimates but fail to reach statistical significance only because of insufficient power to detect small interaction effects (Selvin, 1996, pp. 213–124). Independent samples Student t-tests were conducted to investigate the effect of gender below 200 Hz, at 313 Hz, and at frequencies between 800-1000 Hz for BKBs, based on the data of Cox and Moore (1988), showing effects of gender at these frequencies. Pearson correlations were used to test the association between EHF band levels within subjects across the three speech materials.

## III. RESULTS

Figure 1 shows the average ERB-scaled LTASS calculated for female and male subjects for each speech material (see also Table II in the Appendix). Females consistently had higher EHF levels than males for the BKBs and narratives. For digits, this trend was diminished, particularly for bands ≥15 kHz. The largest gender difference in the EHF region was observed in the 8706-Hz band, with mean differences of approximately 6 dB for narratives, 4 dB for BKBs, and <2 dB for digits. In addition, consistent gender differences were observed for other portions of the spectrum, such as the 806-Hz band, where females had higher levels than males. At the lowest bands (CF ≤123 Hz), males exhibited higher levels than females, owing to lower fundamental frequencies.

**Fig. 1.**
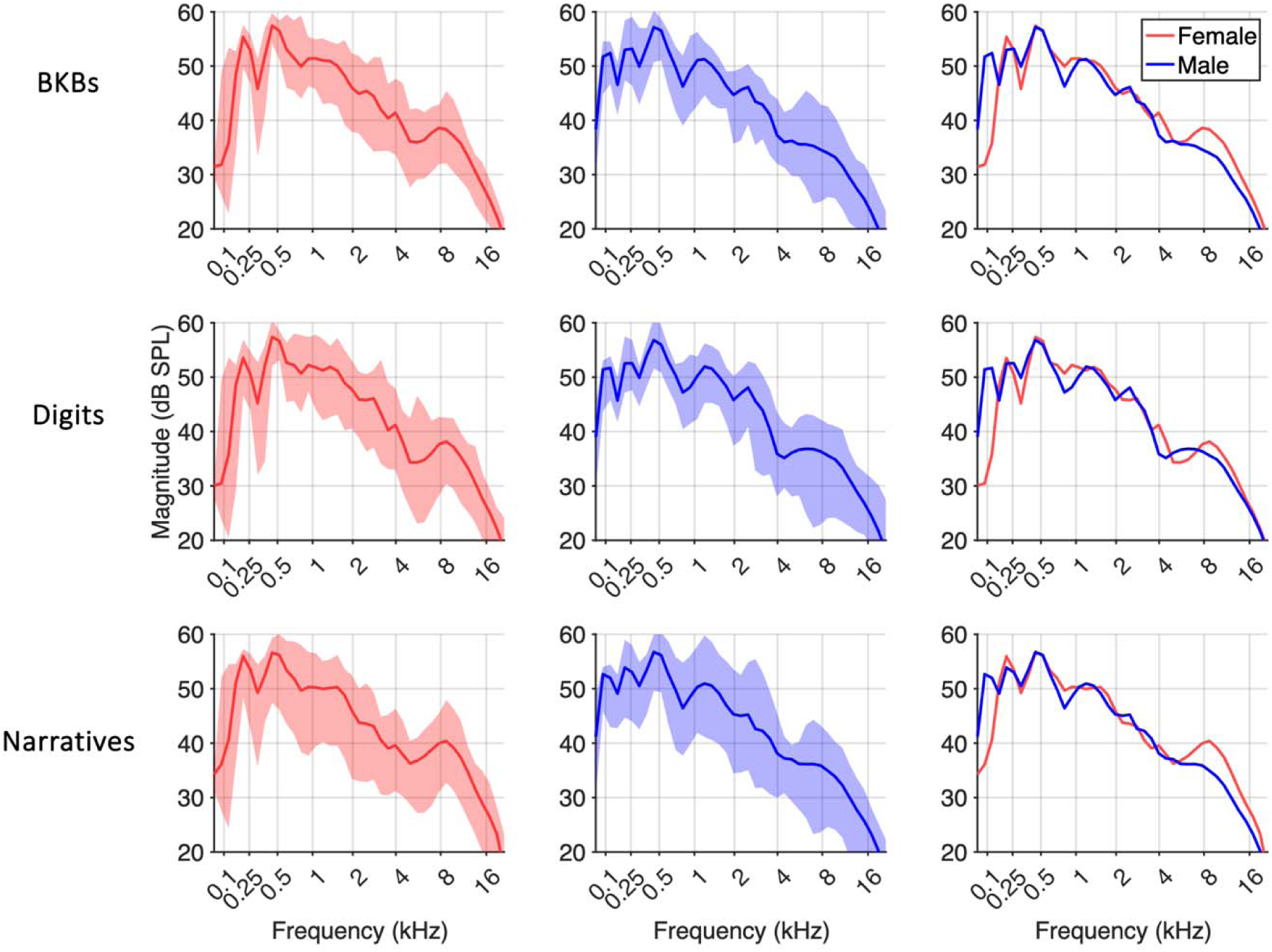
(Color online). Mean ERB LTASS of female and male subjects and their comparison for each speech material on ERB scale. The shaded region shows range (*n* = 15 for each gender for BKBs and digits, and *n* =14 for each gender for narratives).

Figure 2 displays the phonetic content analysis for the three sets of speech material used in this study, along with the phonetic content of the script used in Monson et al. (2012b) and the Rainbow passage for comparison. The phoneme densities for BKBs, narratives, Monson et al. (2012b), and the Rainbow passage were relatively similar. However, digits exhibited some notable differences, including a higher proportion of fricatives and the nasal /n/.

**Fig. 2.**
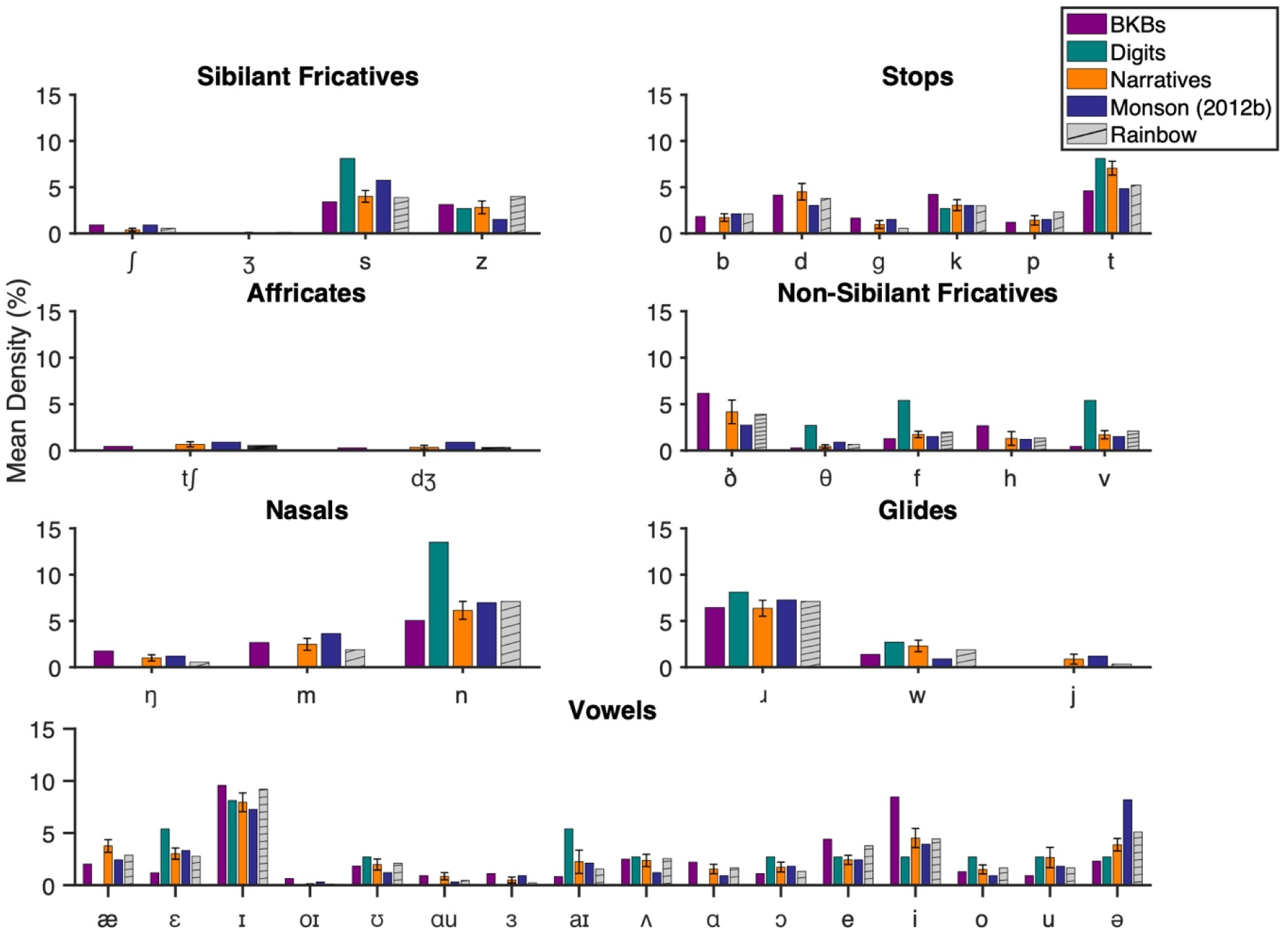
(Color online) Phoneme densities for the three speech materials used in this study. Analyses of the script used in Monson et al. (2012b), and the Rainbow passage were added for comparison. Error bars for narratives indicate standard deviation.

Figure 3 displays comparisons of pairs of speech materials for female and male talkers. Overall, LTASS patterns for all speech materials showed only small differences at EHFs. For female talkers, narratives had the highest EHF levels in general, followed by digits and BKBs. EHF level differences between speech materials were largest (∼2 dB) in the 8706-Hz band but decreased with increasing frequency. Notably consistent differences at other spectral regions include increased energy levels in the 2501-Hz band for digits compared to BKBs and narratives. For male talkers, digits had the highest EHF levels by 1-2 dB, whereas the EHF level differences between narratives and BKBs were less than 0.5 dB.

**Fig. 3.**
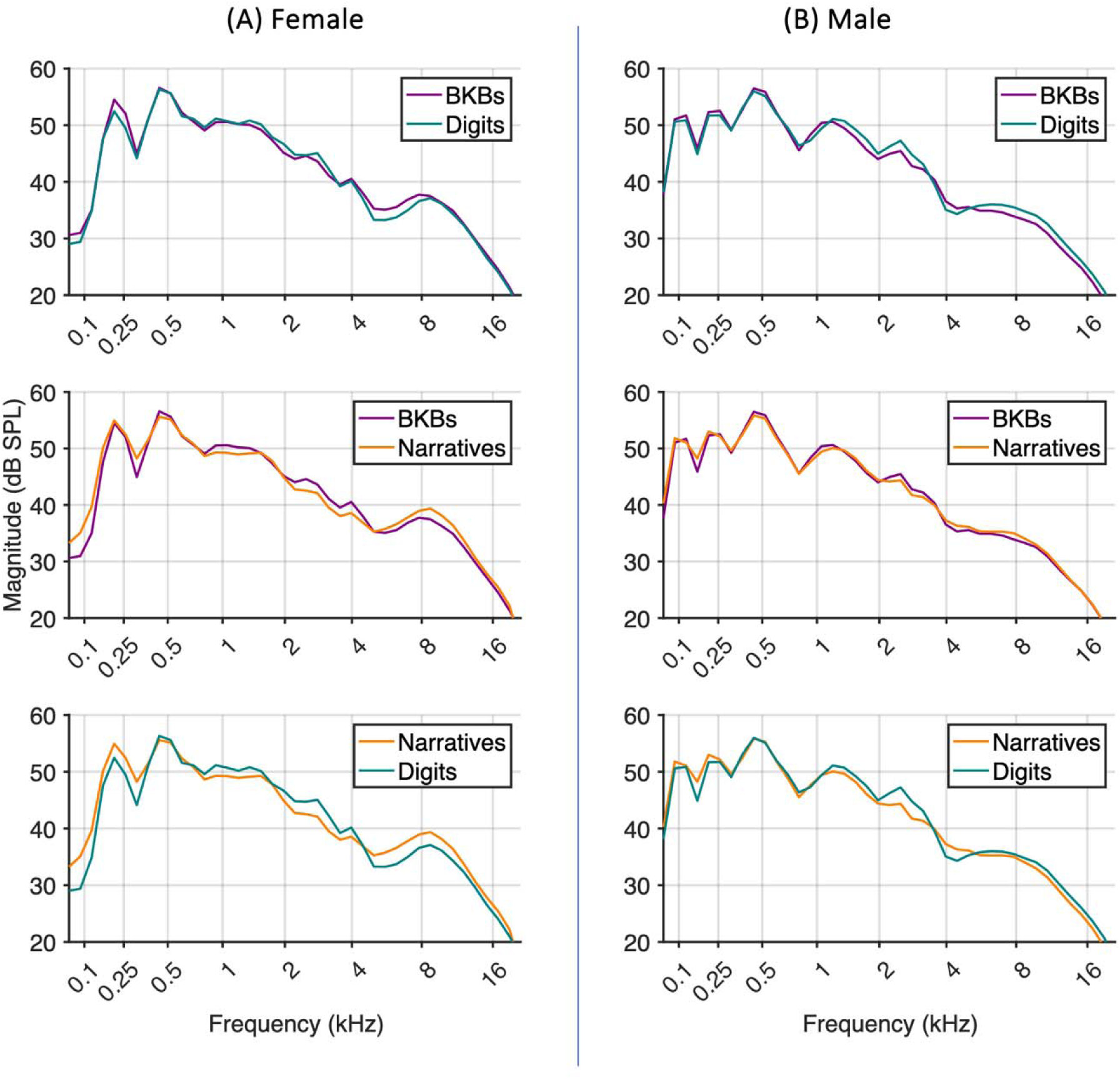
(Color online). (A) Female and (B) male mean ERB LTASS pairwise comparisons for three sets of speech materials on an ERB scale.

These observations were evaluated using a linear mixed-effects model with fixed effects of EHF band, speech material, talker gender, and their two- and three-way interactions. EHF bands (i.e., ERB bands 34-41) were coded as band numbers 0-7 in this model. The reference condition was the female BKBs. The three-way interactions did not approach significance (*p*>0.62) and were removed from the model. The resulting model (Table I) revealed significant main effects and two-way interactions. There was a significant reduction in EHF band level with increasing frequency (i.e. band number). There was a 4.4-dB reduction in EHF level for male talkers, but there was a significant positive interaction between male gender and EHF band, indicating smaller gender differences with increasing frequency. There were also significant interactions between male gender and speech material, indicating larger gender differences for narratives and smaller gender differences for digits, relative to gender differences in BKBs. EHF levels for female narratives were significantly higher (by about 2 dB) than for female BKBs, but the significant male × narratives interaction term indicated this effect was reduced for male talkers. There was no significant difference in EHF level between female BKBs and female digits, but the significant male × digits interaction term indicated this difference was larger for male talkers. A second linear mixed-effects model was computed that included age as factor, however age was not a significant predictor of EHF levels (*p* = .92). Testing for gender differences at the other frequency regions revealed: male talkers had significantly higher levels for ERB bands 3 (CF 87 Hz; t = −12.2, *p* <0.001), 4 (CF 123 Hz; t = −7.6, *p* <0.001), and 8 (CF 313 Hz; t = −3.45, *p* < 0.005), but not ERB band 5 (CF 163 Hz; t = 1.22, *p* =0.23); female talkers had significantly higher levels for ERB bands 14 (CF 806 Hz; t = 3.7, *p* < 0.001) and 15 (CF 924 Hz; t = 2.5, *p* <0.05), in line with previously demonstrated gender effects (Cox and Moore, 1988).

**TABLE I.** Results of the linear mixed-effects model evaluating effects of EHF band, speech material, talker gender, and their interactions on EHF band levels in dB SPL, with overall speech level of 65 dB SPL. The intercept is female BKBs for ERB band 34.

| Term | Estimate | S.E. | t-value | *p*-value |
| --- | --- | --- | --- | --- |
| (Intercept) | 38.75 | 0.8 | 48.53 | <0.001 * |
| EHF band | -2.36 | 0.06 | -37.41 | <0.001 * |
| Male | -4.41 | 1.11 | -3.98 | <0.001 * |
| Digits | -0.31 | 0.37 | -0.83 | 0.4 |
| Narratives | 1.78 | 0.38 | 4.74 | <0.001 * |
| EHF band * Male | 0.35 | 0.06 | 5.56 | <0.001 * |
| EHF band * Digits | -0.02 | 0.08 | -0.23 | 0.82 |
| EHF band * Narratives | -0.14 | 0.08 | -1.75 | 0.08 |
| Male * Digits | 1.85 | 0.35 | 5.24 | <0.001 * |
| Male * Narratives | -0.73 | 0.36 | -2 | 0.04 * |

**TABLE II.**
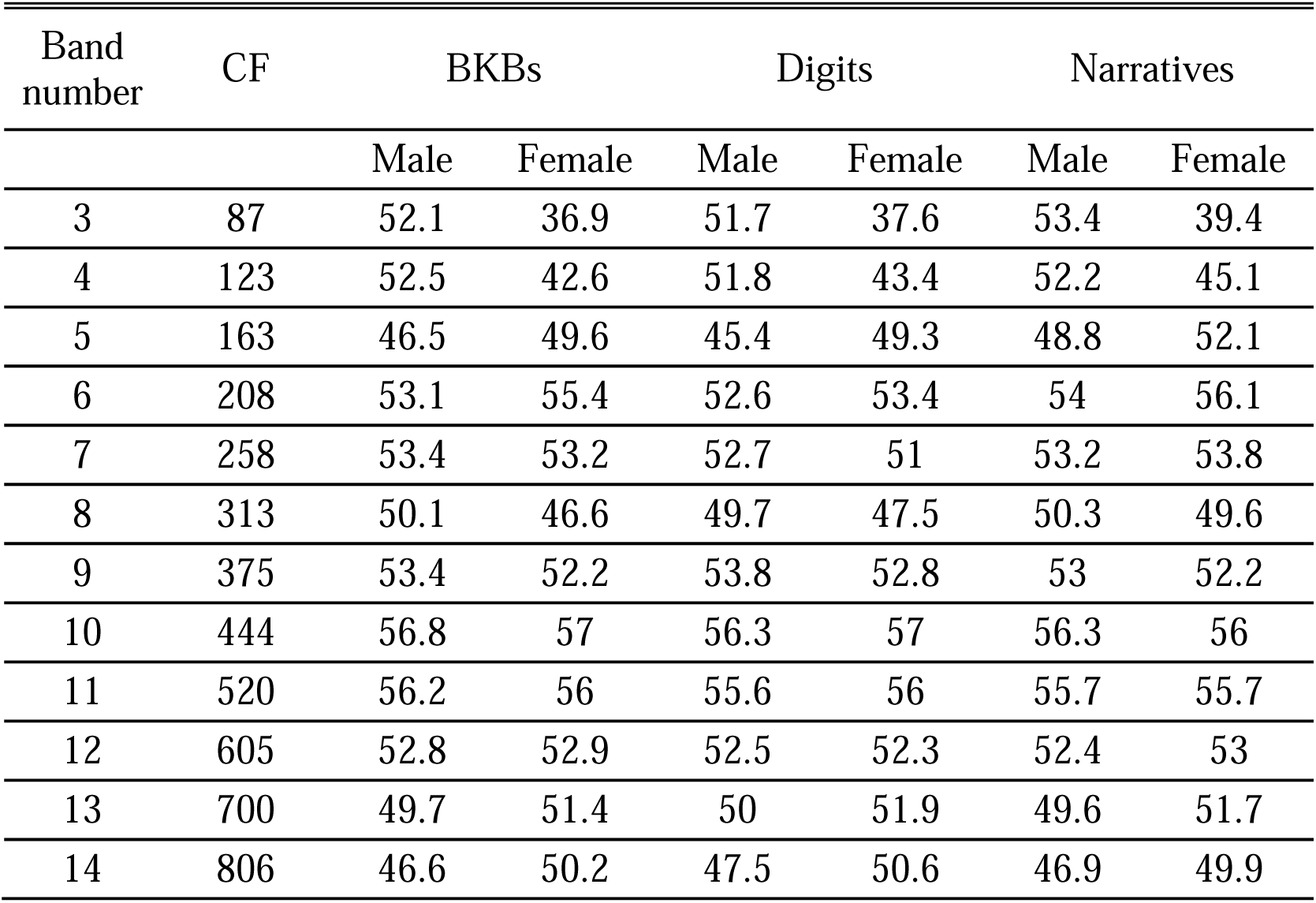
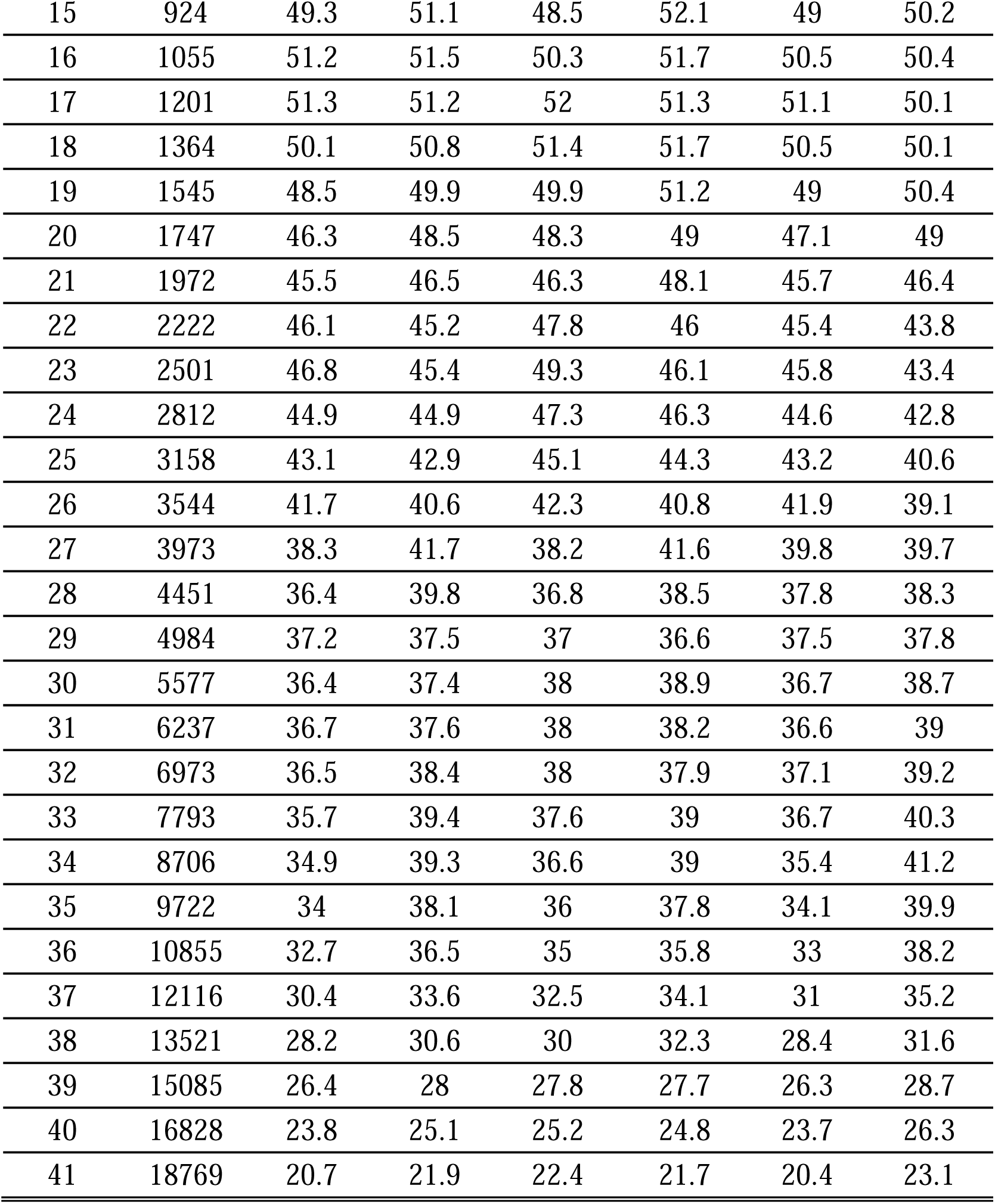
Mean ERB LTASS in dB SPL for each speech material separated by gender. Overall levels are normalized to 65 dB SPL. The center frequencies of each band (CF) are in Hz.

| Band number | CF | BKBs |  | Digits |  | Narratives |  |
| --- | --- | --- | --- | --- | --- | --- | --- |
|  |  | Male | Female | Male | Female | Male | Female |
| 3 | 87 | 52.1 | 36.9 | 51.7 | 37.6 | 53.4 | 39.4 |
| 4 | 123 | 52.5 | 42.6 | 51.8 | 43.4 | 52.2 | 45.1 |
| 5 | 163 | 46.5 | 49.6 | 45.4 | 49.3 | 48.8 | 52.1 |
| 6 | 208 | 53.1 | 55.4 | 52.6 | 53.4 | 54 | 56.1 |
| 7 | 258 | 53.4 | 53.2 | 52.7 | 51 | 53.2 | 53.8 |
| 8 | 313 | 50.1 | 46.6 | 49.7 | 47.5 | 50.3 | 49.6 |
| 9 | 375 | 53.4 | 52.2 | 53.8 | 52.8 | 53 | 52.2 |
| 10 | 444 | 56.8 | 57 | 56.3 | 57 | 56.3 | 56 |
| 11 | 520 | 56.2 | 56 | 55.6 | 56 | 55.7 | 55.7 |
| 12 | 605 | 52.8 | 52.9 | 52.5 | 52.3 | 52.4 | 53 |
| 13 | 700 | 49.7 | 51.4 | 50 | 51.9 | 49.6 | 51.7 |
| 14 | 806 | 46.6 | 50.2 | 47.5 | 50.6 | 46.9 | 49.9 |
| 15 | 924 | 49.3 | 51.1 | 48.5 | 52.1 | 49 | 50.2 |
| 16 | 1055 | 51.2 | 51.5 | 50.3 | 51.7 | 50.5 | 50.4 |
| 17 | 1201 | 51.3 | 51.2 | 52 | 51.3 | 51.1 | 50.1 |
| 18 | 1364 | 50.1 | 50.8 | 51.4 | 51.7 | 50.5 | 50.1 |
| 19 | 1545 | 48.5 | 49.9 | 49.9 | 51.2 | 49 | 50.4 |
| 20 | 1747 | 46.3 | 48.5 | 48.3 | 49 | 47.1 | 49 |
| 21 | 1972 | 45.5 | 46.5 | 46.3 | 48.1 | 45.7 | 46.4 |
| 22 | 2222 | 46.1 | 45.2 | 47.8 | 46 | 45.4 | 43.8 |
| 23 | 2501 | 46.8 | 45.4 | 49.3 | 46.1 | 45.8 | 43.4 |
| 24 | 2812 | 44.9 | 44.9 | 47.3 | 46.3 | 44.6 | 42.8 |
| 25 | 3158 | 43.1 | 42.9 | 45.1 | 44.3 | 43.2 | 40.6 |
| 26 | 3544 | 41.7 | 40.6 | 42.3 | 40.8 | 41.9 | 39.1 |
| 27 | 3973 | 38.3 | 41.7 | 38.2 | 41.6 | 39.8 | 39.7 |
| 28 | 4451 | 36.4 | 39.8 | 36.8 | 38.5 | 37.8 | 38.3 |
| 29 | 4984 | 37.2 | 37.5 | 37 | 36.6 | 37.5 | 37.8 |
| 30 | 5577 | 36.4 | 37.4 | 38 | 38.9 | 36.7 | 38.7 |
| 31 | 6237 | 36.7 | 37.6 | 38 | 38.2 | 36.6 | 39 |
| 32 | 6973 | 36.5 | 38.4 | 38 | 37.9 | 37.1 | 39.2 |
| 33 | 7793 | 35.7 | 39.4 | 37.6 | 39 | 36.7 | 40.3 |
| 34 | 8706 | 34.9 | 39.3 | 36.6 | 39 | 35.4 | 41.2 |
| 35 | 9722 | 34 | 38.1 | 36 | 37.8 | 34.1 | 39.9 |
| 36 | 10855 | 32.7 | 36.5 | 35 | 35.8 | 33 | 38.2 |
| 37 | 12116 | 30.4 | 33.6 | 32.5 | 34.1 | 31 | 35.2 |
| 38 | 13521 | 28.2 | 30.6 | 30 | 32.3 | 28.4 | 31.6 |
| 39 | 15085 | 26.4 | 28 | 27.8 | 27.7 | 26.3 | 28.7 |
| 40 | 16828 | 23.8 | 25.1 | 25.2 | 24.8 | 23.7 | 26.3 |
| 41 | 18769 | 20.7 | 21.9 | 22.4 | 21.7 | 20.4 | 23.1 |

An exploratory analysis was conducted to investigate the correlation for levels in each EHF band across different speech materials. Results showed that individual EHF band levels across speech materials were highly correlated, with strong correlations observed in all bands comparing BKBs vs. narratives (Figure 4). A similar pattern was observed for all bands for the comparison between BKBs and digits, and digits vs. narratives (data not shown).

**Fig. 4.**
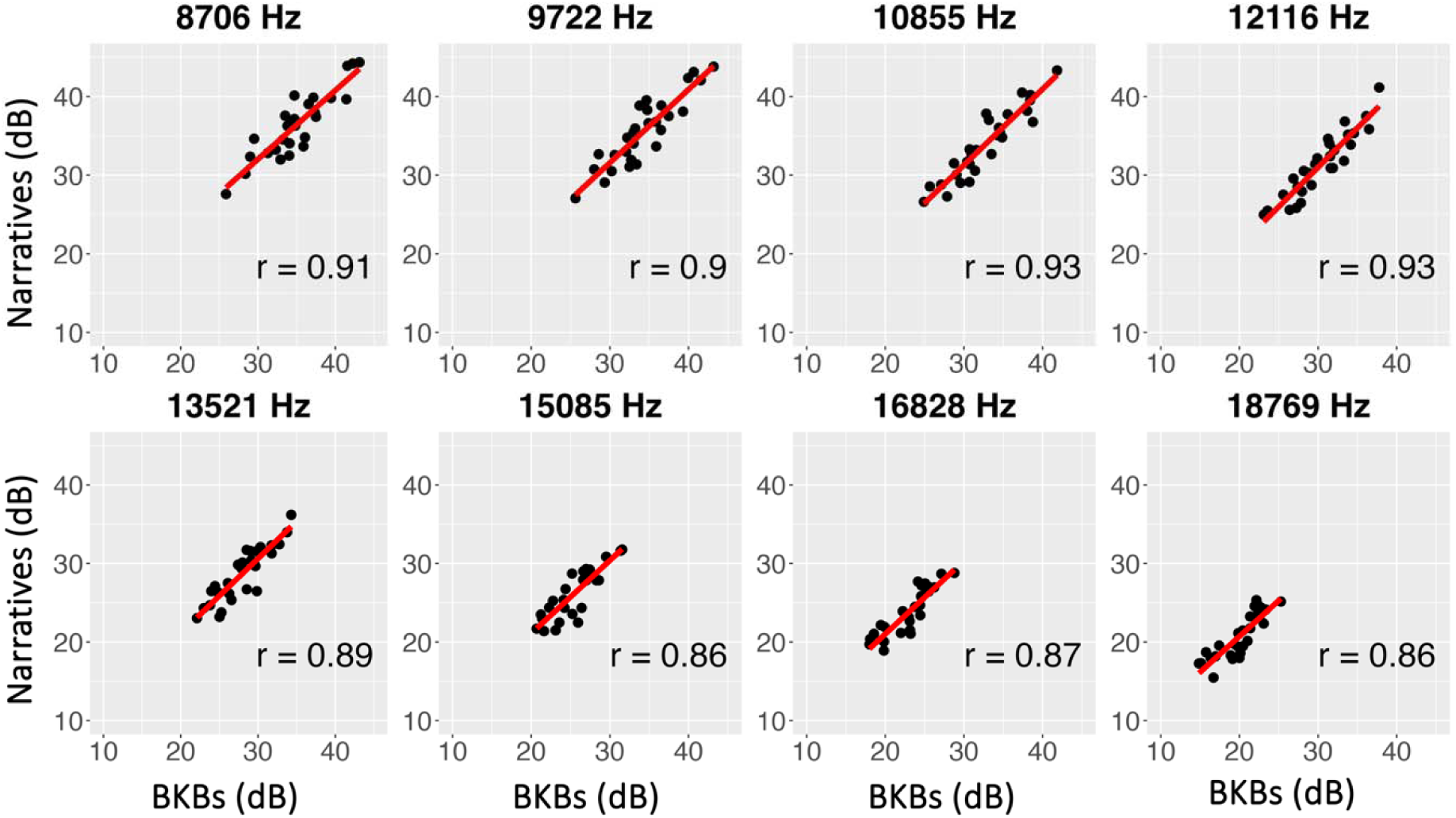
(Color Online). EHF band levels for narratives plotted against BKBs. Each point represents data from one talker. Results for ERBs 34-41 are shown in separate panels, with the associated center frequencies at the top of each panel. The correlation was significant at *p*<0.05 for all bands.

Figure 5 compares third-octave band levels for the mean LTASS for each speech material from the present study with those reported from previous studies (see also Table III in the Appendix). There are some notable differences. For example, LTASS curves from the present study and Monson et al. (2012b) exhibit levels between 1-2 kHz that are consistently ∼4-5 dB higher than levels reported by Moore et al. (2008) and Byrne et al. (1994). LTASS curves from this study and Monson et al. (2012b) also all show a slight increase in band level above 5 kHz which peaks at 8 kHz and decreases thereafter; this pattern was not observed by Moore et al. (2008) or Byrne et al. (1994). All EHF levels in the present study were slightly higher than those reported by Moore et al. (2008) but ∼3-4 dB lower than those reported by Monson et al. (2012b).

**Fig. 5.**
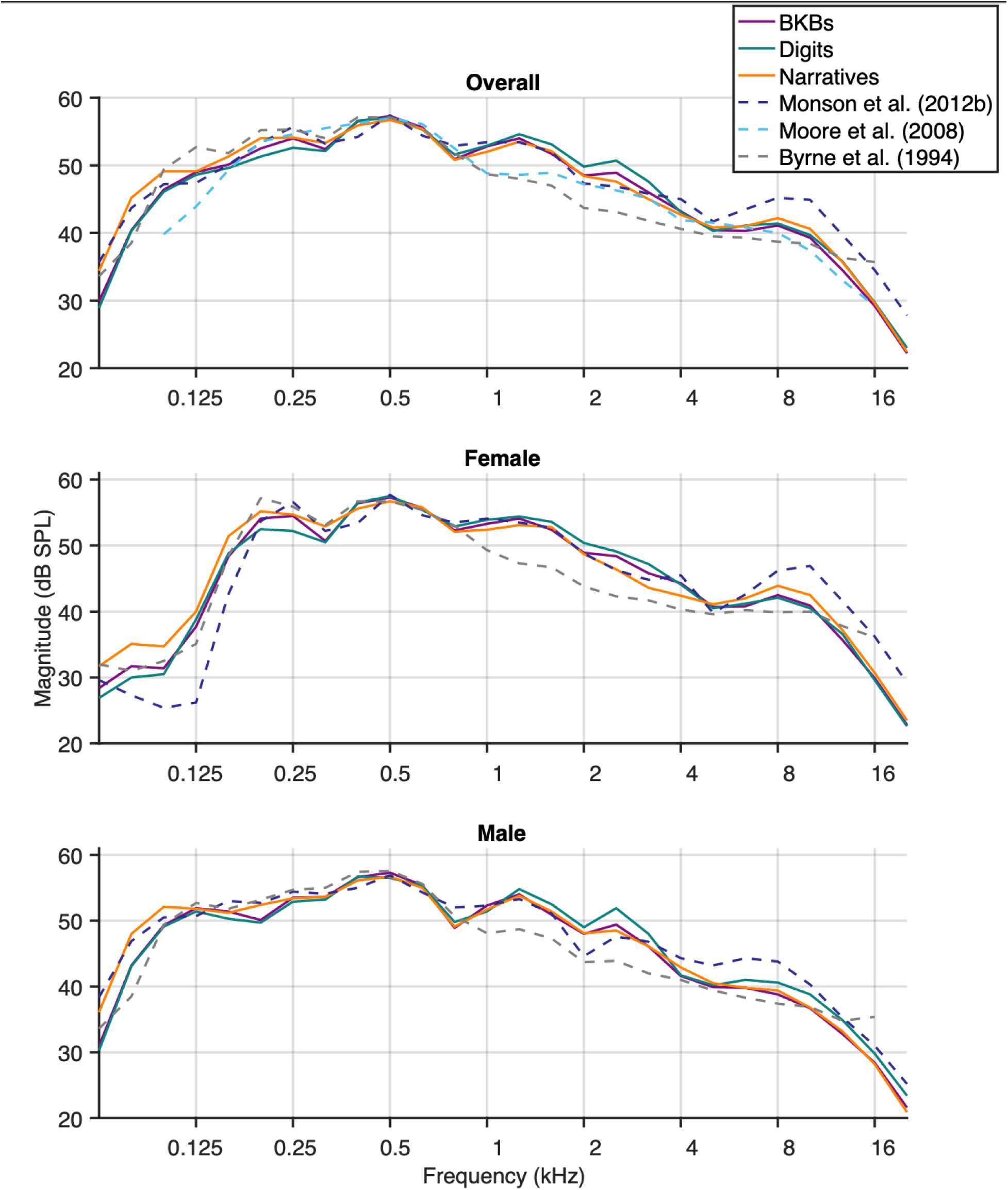
(Color online). Mean LTASS calculated using third-octave bands for overall, male, and female subjects. Solid lines indicate results for the three speech materials evaluated in the present study. Dotted lines show LTASS calculations from Monson et al. (2012b) for semantically unpredictable American English phrases, Moore et al. (2008) for a prose passage in British English, and Byrne et al. (1994) for prose passages collected across 17 different languages and dialects. Levels are set to 65 dB SPL.

**TABLE III.** The mean third-octave band LTASS for the current study and previous studies normalized to an overall level of 65 dB SPL. Center frequencies (CF) reported in Hz and band levels in dB SPL.

| Band CF | BKB | Digits | Narratives | Monson<br>(2012b) | Moore<br>(2008) | Byrne<br>(1994) |
| --- | --- | --- | --- | --- | --- | --- |
| 79 | 40.4 | 40.3 | 45.2 | 43.7 |  | 38.5 |
| 99 | 46.4 | 46.1 | 49.1 | 47.2 | 39.8 | 49.4 |
| 125 | 49 | 48.6 | 49.1 | 47.4 | 43.9 | 52.7 |
| 157 | 50.1 | 49.6 | 51.3 | 50.1 | 49.3 | 51.8 |
| 198 | 52.5 | 51.3 | 54 | 53.3 | 53.5 | 55.2 |
| 250 | 54 | 52.6 | 54.1 | 55.7 | 54.6 | 55.3 |
| 315 | 52.4 | 52.1 | 53.3 | 53.2 | 55.5 | 54 |
| 397 | 56.5 | 56.6 | 55.9 | 54.2 | 56.2 | 57.1 |
| 500 | 57.3 | 57 | 56.7 | 57.4 | 57 | 57.1 |
| 630 | 55.6 | 55.3 | 55.4 | 54.4 | 56.1 | 55.5 |
| 794 | 50.9 | 51.6 | 50.8 | 52.9 | 52.5 | 51.8 |
| 1000 | 52.8 | 52.9 | 52 | 53.4 | 48.8 | 48.7 |
| 1260 | 54 | 54.6 | 53.5 | 53.4 | 48.6 | 48 |
| 1587 | 51.7 | 53.1 | 52.1 | 52 | 48.9 | 47 |
| 2000 | 48.5 | 49.8 | 48.4 | 47.3 | 47.2 | 43.7 |
| 2520 | 48.9 | 50.7 | 47.6 | 46.9 | 46.3 | 43.1 |
| 3175 | 46 | 47.6 | 45 | 45.8 | 45.1 | 41.8 |
| 4000 | 43.2 | 43.1 | 42.7 | 45 | 41.9 | 40.6 |
| 5040 | 40.4 | 40.3 | 40.8 | 41.7 | 41.5 | 39.5 |
| 6350 | 40.3 | 41.1 | 41 | 43.5 | 40.8 | 39.3 |
| 8000 | 41.1 | 41.4 | 42.2 | 45.2 | 40 | 38.7 |
| 10079 | 39.3 | 39.7 | 40.6 | 44.9 | 37.4 | 38.4 |
| 12699 | 34.5 | 35.8 | 35.7 | 39.7 | 33 | 36.3 |
| 16000 | 29.2 | 29.7 | 29.6 | 34.5 | 29.2 | 35.7 |
| 20159 | 22.2 | 23 | 22.4 | 27.8 |  |  |

## IV. DISCUSSION

This study examined the impact of gender and speech material on the LTASS. We utilized three sets of speech materials (BKB sentences, digits, and unscripted narrative) from both male and female talkers. Female speech exhibited significantly higher EHF levels compared to male speech, but this gender difference decreased with increasing frequency and was dependent on speech material. The largest difference was observed in the narratives, where the mean female EHF level was 5.2 dB higher than that for males. For BKBs, this gender difference was 3.8 dB, whereas for digits, it was only 1.7 dB.

It is likely that the gender effect at high frequencies and EHFs is related to anatomical differences between genders that are believed to underlie male-female differences observed in, for example, the peak frequency or spectral mean for voiceless fricatives (Jongman et al., 2000). Because the size of the oral and pharyngeal cavities affects the resonance/peak frequency for voiceless fricatives (Shadle, 1986), smaller cavities associated with the female anatomy may lead to higher peak frequencies and spectral means (Stevens, 1999, pp. 627–628). Since these peak frequencies and spectral means tend to be just below the EHF range (Jongman et al., 2000), these phonemes have relatively high spectral levels at EHFs, which is likely the reason why unvoiced speech tends to provide the dominant contribution to the EHF region of the LTASS (Monson et al., 2011). It has been shown that there are significant gender differences in EHF third-octave band levels for the LTASS of individual voiceless fricatives, particularly /s/ (Monson et al., 2012b). Thus, the gender differences at EHFs observed in the LTASS may be an extension of gender differences in voiceless fricatives, although this effect was smaller in digits which have a high concentration of voiceless fricatives.

We also observed some small effects of speech material on EHF levels that were dependent on gender. We predicted that digits would exhibit higher EHF levels than the other two sets of materials. However, the largest difference in EHF level was between the female BKBs and narratives, with EHF level for narratives only 1.7 dB higher. This BKB vs. narratives difference for males was reduced. Male EHF level for digits was 1.6 dB higher than that for BKBs, but this effect was reduced for females. We expected higher EHF levels for digits because of the higher proportion of voiceless fricatives, which the phoneme distributions indicate.

However, there is also a higher proportion of nasals in the digits (see Figure 2). One spectral feature of nasals is a substantial attenuation in level at high frequencies (Stevens, 1999, pp. 487– 513), which appears to extend into the EHF range (Tabain et al., 2016). Thus, the overrepresentation of nasals in digits may be counteracting the overrepresentation of voiceless fricatives, resulting in little change in LTASS EHF levels. Regardless, despite differences in phoneme distribution and speech material durations, particularly for digits vs. the other materials, the effects of speech material were small relative to the overall gender difference of ∼4 dB at EHFs.

It is intriguing that the LTASS differences at EHFs were not consistent across genders. One possible explanation for this is that gender differences in other factors known to affect the LTASS (e.g., emotion, prosody; Williams & Stevens, 2005) may have been more pronounced for natural narrative speech than for the scripted speech, which could have had a gender-specific effect on the narratives LTASS levels (Přibil and Přibilová, 2012; Williams and Stevens, 2005). Another possibility that cannot be ruled out is that the audio prompt (female voice for BKBs, male voice for digits) may have influenced the talkers’ production, including at EHFs. Additionally, we found that EHF levels were highly correlated across different speech materials. This finding suggests relative consistency in EHF level for a given talker uttering different speech materials.

Comparing our data to previously published data, we replicated reported gender effects at both ends of the speech spectrum: <163 Hz and >8 kHz (Cox and Moore, 1988). Replicating the findings of Monson et al. (2012b) for gender differences at EHFs suggests this previous finding was not specific to the young, vocally trained population used in that study. We also replicated gender differences previously observed at 800-1000 Hz (Cox and Moore, 1988; Monson et al., 2012b), where female levels in the present study were ∼3 dB higher than male levels, and at 315 Hz, where the male level was 3.5 dB higher than the female level. The LTASS data for speech materials analyzed here align very well with the Monson et al. (2012b) data up to approximately 5 kHz (Fig. 5; Table III). Although the speech material differed for Monson et al. (2012b), our analysis showed that the phonetic distribution for Monson et al. (2012b) did not differ substantially from the BKBs and narratives (Figure 2). The similarity between the phonetic distributions, along with other similarities in the approach (Class I microphones, anechoic conditions), may have led to these similarities in the LTASS data.

However, there were also some discrepancies between our data and previously published data. For example, Monson et al. (2012b) reported EHF levels that were 3-4 dB higher than those measured in the present study. EHF levels measured here were more similar to, but slightly greater than, those reported by Moore et al. (2008). The use of different speech materials may have contributed to these differences, although our data suggest the effect of speech material on EHF levels would be smaller than the discrepancy between the present data and those of Monson et al. (2012b). Both of these studies used Class I precision microphones located at the level of the mouth, but differences in study parameters included: recording distance (60 cm versus 1 m), recording posture (standing versus seated), and participant demographics (young, vocally trained talkers versus diverse talkers) in the Monson et al. (2012b) study compared to present study, respectively. It may be that these factors influenced EHF levels. For example, it is possible that vocally trained singers produce more EHFs than the typical population. Additionally, Monson et al. (2012) used low-predictability sentences with alternating syllabic strength (e.g., amend the slower page). This type of material could introduce differences in speech rate and prosodic structure (Smiljanić and Bradlow, 2009) that could possibly have an effect on the the LTASS at EHFs. In the present study, there was substantial individual variability in levels at EHF bands, with a spread of 10-15 dB, depending on frequency. Given this variability, it is feasible that EHF level differences across studies could be attributed to talker variability rather than the speech material or other recording parameters. Finally, whereas Monson et al. (2012b) LTASS data align well with the present data at frequencies ≤5 kHz, Moore et al. (2008) and Byrne et al. (1994) LTASS data show relatively large differences at low (100-200 Hz) and middle (1-4 kHz) frequencies. These differences could arise from differences in recording setup, but may also be related to language differences in recordings. Byrne et al. (1994) calculated a composite LTASS using 17 languages and dialects, and the Moore et al. (2008) recordings were British English, whereas the recordings analyzed here and in Monson et al. (2012b) were American English.

One question that arises from these data is whether these EHF level differences have perceptual consequences. A caveat in this discussion point is that there are many additional real-world factors not assessed here that will affect the EHF cues a listener actually receives, including hearing loss, talker head orientation, listener head orientation, and distance. However, a 4-dB increase in EHF levels could theoretically improve audibility of EHF cues for female speech relative to male speech. Most studies that have reported effects of EHF cues for speech perception have used female speech (Lippmann, 1996; Monson et al., 2019; Motlagh Zadeh et al., 2019; Polspoel et al., 2022; Stelmachowicz et al., 2007; Trine and Monson, 2020), demonstrating that EHF cues provide a robust benefit for female speech recognition in noise for normal-hearing listeners. A few studies have examined male speech recognition and EHFs for normal-hearing listeners (Levy et al., 2015; Polspoel et al., 2022; Stelmachowicz et al., 2001), but only one of these studies used full-band (20-kHz bandwidth) stimuli. Polspoel et al. (2022) observed an EHF benefit for male digits-in-noise recognition, showing better performance when using full-band stimuli as compared to stimuli that were low-pass filtered at 8 kHz. Levy et al. (2015) found no significant difference in male speech recognition between 8-kHz and 10-kHz bandlimited speech. Stelmachowicz et al. (2001) demonstrated that an increasingly wider bandwidth (out to 9 kHz) was beneficial for female speech recognition, whereas male speech recognition performance did not improve appreciably when bandwidth increased from 5 kHz to 9 kHz. Whether increasing bandwidth beyond 9 kHz would have improved speech recognition for that study is unknown. Thus it is not clear to what extent providing the full complement of EHF cues contributes to male speech recognition as the only study to assess full-band (20-kHz bandwidth) male speech recognition and EHF cues was Polspoel et al. (2022). For detection of EHFs in speech, Monson et al. (2014b) found no effect of talker gender, but Monson & Caravello (2019) found a small effect of gender, with listeners displaying slightly better detection thresholds for EHFs in female speech. Additional investigation of the effect of talker gender differences on the EHF benefit for speech recognition is warranted.

## ACKNOWLEDGMENTS

We thank our study participants. This project was supported by NIH grant number R01-DC019745.

## AUTHOR DECLARATIONS

### Ethics Approval

Informed consent was obtained from all participants and all data collection procedures were approved by the Institutional Review Board at Boys Town National Research Hospital.

## DATA AVAILABILITY

The data supporting the findings of this study are available upon reasonable request.

## APPENDIX A: MEAN LTASS LEVELS

